# A quantum state of mitochondria in the living cell

**DOI:** 10.64898/2026.09.10.750628

**Authors:** Yu Yang, Zhenglong Gu, Bo Song

## Abstract

The high energy-efficiency of life is hard to understand only with classical physics. Many efforts have been made to study its mechanism based on quantum mechanics; the progress is nevertheless slow due to lack of experimental evidence with living cells. Here, combining experiments on cells, tissues and mitochondria with a theoretical model, we demonstrate a quantum state of mitochondria, which can be employed to modulate ATP production in living cells. We found an anomalous 71.0-THz oscillation mode only in living cells and tissues, which is highly determined by intact structure of mitochondria, and cannot be assigned to any specific molecules. Based on experimental data, a quantum model of light-matter coupling was introduced to trace the origin of this mode. Our calculations suggest a quantum superposition state of functional mitochondrion that forms by the coupling of light and lipid CH_2_ bonds in functional cristae, and induces a splitting of the intrinsic CH_2_ vibration mode of 87 THz to two levels at 71 THz and 103 THz, respectively. The former can be observed only in living cells and tissues; whereas the latter falls in the range (90–110 THz) of biomolecular and water vibrations, thus indistinguishable. Additional experiments revealed this mitochondrial quantum state able to serve as an efficient channel to modulate ATP production. Our findings provide a quantum mechanics view for understanding living cells, and it will be interesting to further explore whether such quantum state could act as a channel for energy metabolism, and even information transmission in life.

## Introduction

Life utilizes energy with a high efficiency (e.g., an adult human operates with a power of ∼100 W, whereas ∼20 W for the brain) [1,2]. Classical physics cannot explain this phenomenon, and many efforts have been made to study the underlying mechanism from quantum mechanics [3–8]. For explaining the high efficiency of human brain, Penrose suggested a model of quantum coherence in neural microtubules [3]; molecular dynamics simulations together with analyses of de Broglie wave lengths demonstrated a quantum coherent oscillation in neural ion channels [4]. As to the harvesting of visual and near-infrared light in plants, quantum coherence-based energy transfer was proposed [5,6], and Leggett et al. indicated that the polariton of light-exciton coupling is able to explain the photosynthetic ultra-efficiency [7]. Recently, evidence for nonthermal effects of mid-infrared (MIR) light has been discovered in multiple biological systems. MIR stimulation with a specific frequency was shown to have resonant interactions with ion channels, exerting nonthermal and energy-efficient modulation on neuronal signaling [9,10]. MIR photons associated with ATP hydrolysis were reported to resonantly influence DNA replication and assembly in model systems [11–13]. Interestingly, our theoretical work suggested that 87-THz MIR light has a capability to couple with myelin sheath, causing a quantum state, and playing an important role in neural communications [14–16]. Despite these progresses, experimental evidence for biological quantum states, especially at the cellular level as a key to understanding the high energy-efficiency of life, remains lacking.

The emergence of mitochondria represented an essential leap in energetic evolution of life [17,18]. Mitochondria as powerhouses of eukaryotic cells generate most ATP in body, and their dysfunction is associated with aging and pathologies [19–22]. The mitochondrial energy transduction efficiency can reach ∼90%, already beyond the conventional framework of thermodynamics [23,24]. Electron tunneling in respiratory chain complexes of mitochondria was proposed to play an important role in ATP production [25]. Lee et al. developed a method based on biological electron tunneling junctions, and explored the electron transfer dynamics in mitochondrial cytochrome *c* [26]. Friedrich et al. revealed that in the respiratory complex I, the iron-sulfur chain regulates the electron tunneling rate, enabling efficient energy conversion [27].

Nevertheless, few experiments are conducted at the cellular level, limiting our understanding on quantum dynamics of energy efficiency in living systems. In this study, an anomalous oscillation mode of 71.0 THz was observed only in living cells and tissues, which is highly determined by functional mitochondria, but cannot be assigned to any specific molecules. Our experiment data-based analyses and theoretical calculations indicated that a quantum state of mitochondrion forms by the coupling of light and lipid CH_2_ bonds in functional cristae, corresponding to the mitochondrion-related oscillation mode we observed. Further experiments revealed that the mitochondrial quantum state provides an energy-efficient channel to modulate ATP production in living cells.

## Results and Discussion

### 1. Mitochondrion-Related 71.0-THz Oscillation Mode in Living Cells and Tissues

Characteristic peaks in spectra of quantum systems are closely associated with the fundamental concepts (e.g., energy levels, oscillation modes) of quantum mechanics, and even applied to analyze the property of quantum superposition [28]. Therefore, to explore the quantum oscillation mode (i.e., quantum state) in living systems, we first performed MIR spectral analyses on the commonly-used human renal epithelial cell line HEK-293T and mouse kidney tissue, with ground dry samples as controls. The living and dry samples were measured by the Fourier transform infrared (FTIR) spectroscopy with the modes of attenuated total reflectance (ATR) [29,30] and transmission [31], respectively.

There is a group of characteristic peaks in the 68–72 THz range of FTIR spectra, only occurring in the living cells and tissue, but not the ground dry ones. We observed three groups of characteristic peaks around 50 THz, 70 THz and 87 THz, respectively (Fig. 1a,d). The first and third groups took place both in the living and ground samples, and agreed with the well-known vibration modes of peptide amide I and CH_2_ (& CH_3_) stretch, respectively [31]. Surprisingly, the second group around 70 THz in the range of 68–72 THz only occurred in the living samples, but not in the ground ones, while according to the reported spectra of biomolecules, there should not exist any characteristic peaks in the range of 60–80 THz (Supplementary Fig. 1) [31].

**Fig. 1.**
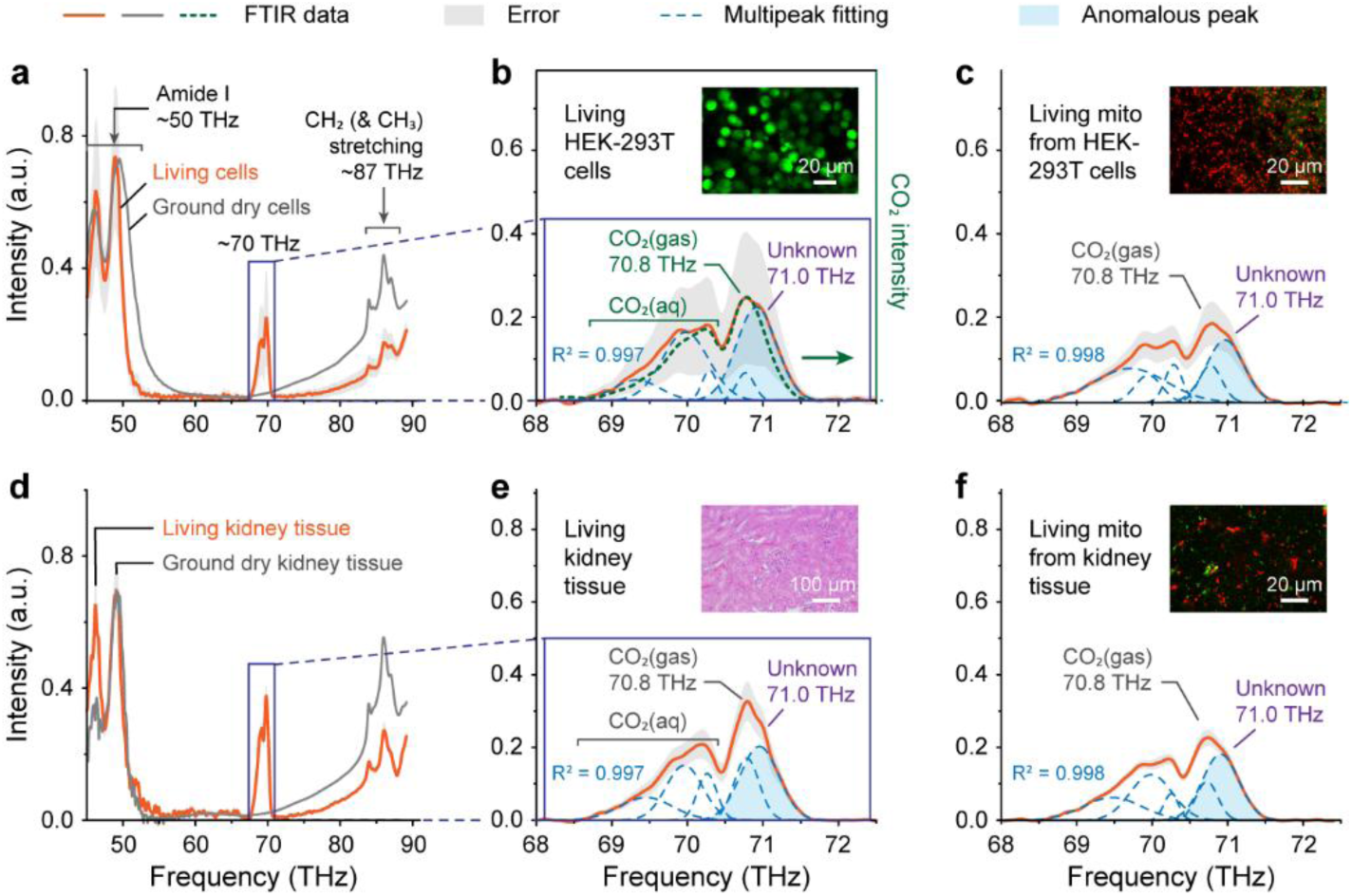
| Mitochondrion-related characteristic peaks at a frequency of 71.0 THz only in spectra of living cells and tissue. **a-c)** HEK-293T cells (**a**), zoom-in of the 68–72 THz range (**b**) and living mitochondria extracted from HEK-293T cells (**c**). In Panel **b**: The green dotted curve indicates the FTIR spectrum of carbon dioxide; CO_2_(aq) and CO_2_(gas) stand for the dissolved and gaseous states of CO_2_ molecules, respectively. **d-f)** Kidney tissue of mice (**d**), zoom-in of the 68–72 THz range (**e**) and mitochondria extracted from kidney tissue (**f**). Inset: Micrograph (calcein AM staining for living cells, H&E for tissue; JC-1 staining for mitochondria with higher (red) and lower (green) membrane potentials). The tangerine and gray curves indicate the FTIR-measured data, with the gray shadow for the error. The blue dashed curve denotes the multipeak fitting (the determination coefficient R^2^ > 0.995); the light-blue shadow means the mitochondrion-related characteristic peak.

To study the surprising peaks observed in the 68–72 THz range, we performed fine analyses on the spectra of living HEK-293T cells and kidney tissue. There were two groups of characteristic peaks in 68.5–70.4 THz and 70.4–71.6 THz, respectively (Fig. 1b,e). The multipeak analysis with the determination coefficient R^2^ > 0.995 showed that the first group consisted of at least three peaks, and the second contained a main peak at 70.8 THz and a shoulder peak at 71.0 THz. It is well known that the MIR range of 68–72 THz is closely related with the molecular vibration of carbon dioxide [32]. We thus measured the spectrum of this important metabolite, and observed two peak groups in 68.5–70.4 THz and at 70.8 THz, respectively, corresponding to the fingerprint peaks of dissolved CO_2_ (labelled CO_2_(aq)) and gaseous CO_2_ (labelled CO_2_(gas)) (Fig. 1b, Supplementary Fig. 2) [32]. Thereby, the first group of peaks in the 68.5–70.4 THz range of living sample spectra was assigned to the vibration of CO_2_(aq), and the second group’s main peak at 70.8 THz was assigned to the CO_2_(gas) vibration. However, the origin of the second group’s shoulder peak at 71.0 THz was still unclear. Interestingly, the isolated mitochondria showed similar patterns in the spectrum (Fig. 1c,f). Therefore, the unknown characteristic peak at 71.0 THz in the spectra of living HEK-293T cells and kidney tissue is closely associated with a certain oscillation mode of the mitochondria rather than the vibrations of any specific molecules.

Next, if the 71.0-THz peak relates to the mitochondrial oscillation, it should widely exist in various living systems. We thus further investigated MIR spectra of mouse liver, heart and skeletal muscle tissues together with the mitochondria extracted from them. For the living liver tissue and its isolated mitochondria, two groups of characteristic peaks were observed in the ranges of 68.5–70.4 THz and 70.4–71.6 THz, respectively (Fig. 2a). Multipeak analysis (R^2^ = 0.997) showed that the first group consisted of at least three peaks, assigned to the vibration of CO_2_(aq). The second group contained a main peak at 70.8 THz and a shoulder peak at 71.0 THz, where the main peak was assigned to the CO_2_(gas) vibration and the shoulder remained unknown. The similar patterns also occurred in the spectra of the heart, skeletal muscle tissues and their isolated mitochondria (Fig. 2b,c). These results indicate that the mitochondrion-related anomalous oscillation mode of 71.0 THz exists in the living liver, heart and skeletal muscle tissues as well.

**Fig. 2.**
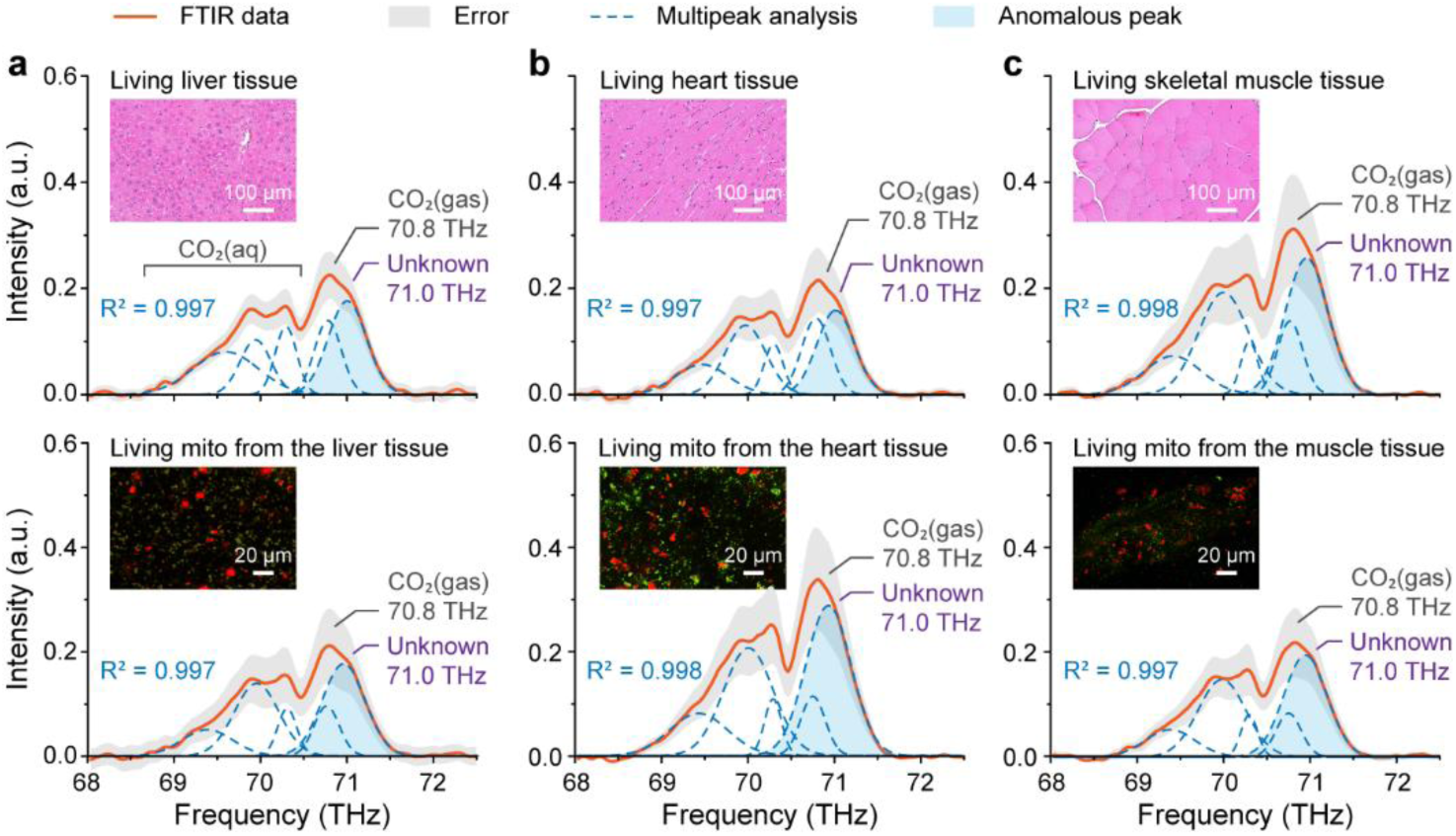
| Mitochondrion-related characteristic peaks at a frequency of 71.0 THz in the spectra of living liver (a), heart (b) and skeletal muscle (c) tissues of mice together with their isolated mitochondria. Upper: Tissue. Lower: Mitochondria from the tissue. Inset: Micrograph (H&E staining for tissue, JC-1 staining for mitochondria with higher (red) and lower (green) membrane potential). The tangerine solid and blue dashed curves mean the FTIR-measured data and multipeak fitting (the determination coefficient R^2^ > 0.995), respectively; the gray shadow denotes the error. The light- blue shadow indicates the mitochondrion-related characteristic peak.

Overall, we observed an anomalous characteristic peak at 71.0 THz in multiple kinds of living samples, including HEK-293T cells, and mouse tissues of kidney, liver, heart and skeletal muscle, which is closely related with a certain oscillation mode of the mitochondria rather than the vibrations of any specific molecules. This anomalous peak disappears when the sample structure is destroyed, reflecting that the corresponding oscillation mode is highly determined by the mitochondrial structure.

### 2. Origin of the Mitochondrion-Related Oscillation Mode

To explore the origin of the mitochondrion-related 71.0-THz mode, through analyzing morphological, structural and optical characteristics of functional mitochondria, we built up a quantum model of light- mitochondrion coupling, and compared its properties with those of the above 71.0-THz mode.

#### Characterized morphology of mitochondria in the living HEK-293T cells

Well-grown HEK-293T cells, labelled with fluorescent dyes Hoechst-33342 (nuclei) and JC-1 (mitochondria) [33,34], were employed for three-dimensional (3D) confocal fluorescence microscopy. The average of measured nuclear diameters was 11.2 ± 1.4 μm (n = 30 nuclei) (Supplementary Fig. 3a,b), consistent with the previously reported value of 11.6 ± 0.4 μm [35]. With JC-1, the mitochondria of higher and lower membrane potentials (MMP) exhibit red (JC-1 aggregates) and green (JC-1 monomers), respectively, corresponding to the cases with higher and lower energy metabolism activities, i.e., energy-active and -inactive mitochondria [34]. In our experiments, these two kinds of mitochondria were observed to separately distribute in different regions, and most of the energy-active ones tended to align along the z-axis (i.e., normal of confocal dish plane) (Fig. 3a).

**Fig. 3.**
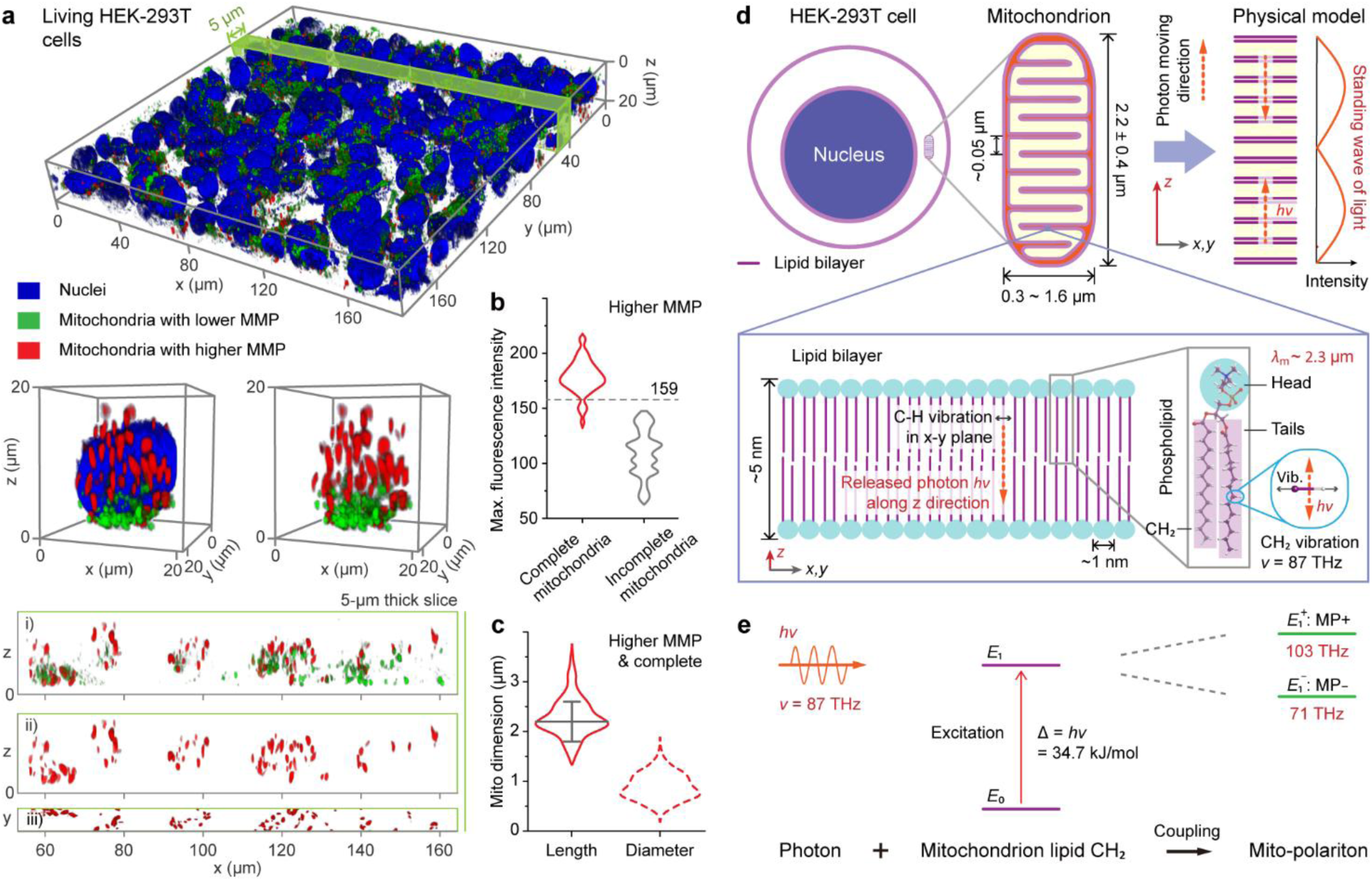
| A 71-THz quantum state of living HEK-293T cellular mitochondria. **a-c)** Mitochondrial morphology. **a)** Confocal fluorescence micrographs of living HEK-293T cells. The blue, green and red denote the nucleus (Hoechst 33342 stained), mitochondria (JC-1 stained) with lower and higher mitochondrial membrane potentials (MMP), respectively. Top and middle: Three-dimensional images. The green ribbon means the cropped example of 5-μm thickness. Bottom: Typical cropped example of mitochondria in the x-z (i, ii) and x-y (iii) views. **b)** Maximum-fluorescence intensity of complete (red) and incomplete (gray, due to cutting boundaries, seeing x-y view of cropped example) higher- MMP mitochondria. **c)** Length (solid) and maximum-diameter (dashed) of complete higher-MMP mitochondria. **d-e)** Quantum state of HEK-293T cellular mitochondrion. **d)** Multi-scalar analyses on mitochondrial structures. Upper left, upper middle and lower: cell, mitochondrion and lipid bilayer. The label *λ*_m_ indicates the wavelength of 87-THz light in the mitochondrion; *ν* = 87 THz denotes the frequency of CH_2_ bond. Upper right: A simplified model of mitochondrion and 87-THz light. **e)** Mito- polariton (MP) caused by coupling of 87-THz light with CH_2_ bonds in the lipid bilayer of mitochondrial cristae as well as their quantized energy levels. *E*_0_ and *E*_1_ stand for the ground and excited states of mitochondrial lipid CH_2_ vibration, respectively; 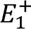 and 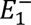 for the split levels of *E*_1_ caused by the coupling of light, labelled MP+ and MP−, respectively.

To facilitate fine analyses on the morphology of energy-active mitochondria, we cropped the 3D sample image along the y axis. The typical slices of sample in paired x-z and x-y views are shown in Fig. 3a lower (more in Supplementary Fig. 3c). The small light-red patterns in the x-z image mean the incomplete mitochondria caused by splicing, and others for the complete. Maximum fluorescence intensities (MFIs) of the incomplete and complete mitochondria were distributed in the ranges of 63– 147 and 132–217, respectively, with a little overlap between them (Fig. 3b). A critical value of 159 was thus introduced to filter out all the incomplete mitochondria. The measured lengths (*L*_m_) of energy-active complete mitochondria with MFIs > 159 were normally distributed in 1.3–3.7 μm, with an average of 2.2 ± 0.4 μm (n = 100) (Fig. 3c). Their maximum diameters (*d*_m_) were non-normally in 0.3–1.6 μm, consistent with the well-known features (0.5–1.2 μm) [36]. These results indicate that the lengths of functional mitochondria in HEK-293T cells are concentrated at 2.2 μm with a slight fluctuation, and the diameters are smaller than 1.6 μm.

#### Structural and optical characteristics of energy-active mitochondria

The structural property of functional mitochondria was multi-scalarly analyzed (Fig. 3d). The length *L*_m_ of mitochondria was 2.2 ± 0.4 μm, and the diameters *d*_m_ were distributed in 0.3–1.6 μm significantly smaller than *L*_m_, reflecting a structure of micron rod. A mitochondrion mainly consists of mitochondrial cristae as perpendicular to the mitochondrial axis, i.e., parallel to each other, with an inter-cristae distance of ∼0.05 μm much smaller than the mitochondrial length (Fig. 3d upper) [37]. The cristae are composed of a lipid bilayer as consists of phospholipid with a head and two tails (Fig. 3d lower). There are ∼30 of CH_2_ bonds in the two tails above. All the lipid CH_2_ bonds tend to be parallel to the lipid bilayer plane, with a huge area density of ∼8 × 10^11^ μm^-2^, and a much smaller inter-bond distance (10^-4^ μm level) [38] than the mitochondrial diameter (0.3–1.6 μm). Based on the inter-cristae distance (∼0.05 μm), the bulk density of CH_2_ bonds was estimated to be 10^13^ μm^-3^, also high extremely. These analyses indicate that all cristae CH_2_ bonds in an energy-active mitochondrion form a micron rod with an ordered high-density structure, well consistent with the previously reported higher density and higher order of cristae in functional mitochondria [39].

Next, we studied the light-related property of energy-active mitochondria. The frequency of CH_2_ stretch mode was around 87 THz (Fig. 1a), and thus a lipid CH_2_ bond can be resonantly excited by 87-THz light. In principle, the excited CH_2_ bond will emit 87-THz photons when relaxing. Due to the structural character of ordered high-density CH_2_ bonds in an energy-active mitochondrion, the above emitted photons mainly propagate along the mitochondrial axis, and then are most likely absorbed by another CH_2_ bond with resonance (Fig. 3d). This exchange of photons between the cristae CH_2_ vibrations will cause a resonant coupling of the light and vibrations, which strength highly depends on the density and order of CH_2_ bonds, potentially correlated with the dependence of mitochondrial energy metabolism activity on the cristae’s density and order [39].

As for the spatial property of 87-THz light in the mitochondrion, its vacuum wavelength (*λ*_0_) is 3.4 μm, and the refractive index (*n*) is ∼1.5 for a lipid bilayer-rich structure of living systems [40]. The wavelength (*λ*_m_) in mitochondria was thus determined to be ∼2.3 μm by *λ*_m_ = *λ*_0_/*n*, consistent with the mitochondrial length *L*_m_ (2.2 ± 0.4 μm) of living HEK-293T cells. This dimension matching enables a standing wave mode of 87-THz light (Fig. 3d upper right), and then a cavity-confinement effect of the mitochondrion on the light with help of the resonant coupling between light and lipid CH_2_ bonds in cristae.

#### A quantum model built up for the energy-active mitochondria

Based on the analyses above of morphological, structural and optical characteristics, we introduced a lipid bilayer-composed cavity model for the light-matter coupling system of energy-active mitochondrion (Fig. 3d upper right). The cavity length was set to 2.3 μm (i.e., *λ*_m_), and the light of propagating along the cavity axis (i.e., *z* direction) was majorly considered. The Hamiltonian of this model was written as,

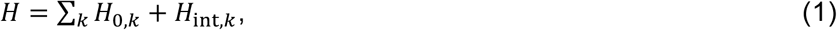

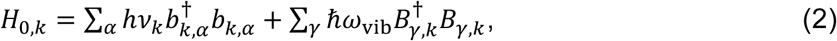

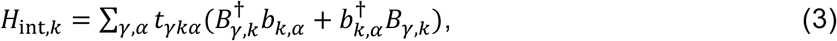

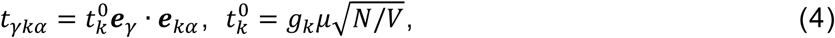

where *b*^†^ (or *b*) and *B*^†^ (or *B*) indicate the creation (or annihilation) operators of mitochondrion cavity- confined photon and lipid CH_2_ vibron, respectively. The labels ℏ*ω*_vib_, *hν_k_* and *t_γkα_* denote the energies of CH_2_ vibron, photon and their coupling, respectively, with *ω*_vib_ = 2π*ν*_vib_ and ℏ = *h*/2π, where *h* = 6.63 × 10^-34^ J s = 0.4 kJ mol^-1^ THz^-1^ represents the Planck constant; *ν*_vib_ and *ν_k_* mean the frequencies of CH_2_ vibration and photon, respectively. The wavevector *k* was along the *z* direction, i.e., *k* = *k_z_*. The label ***e****_kα_* indicates the unit vector of light polarization direction *α* = 1 and 2; *μ* and ***e****_γ_* stand for the magnitude and direction of the *γ*-th CH_2_ dipole vector **μ***_γ_* = *μ**e**_γ_* in the lipid bilayers, respectively, and *g_k_* = (*hv_k_*/2*ε*_m,0_)^1/2^ with *ε*_m,0_ for the mitochondrial permittivity excluding the contribution from lipid CH_2_ vibrons. *N* and *V* indicate the CH_2_ bond number and the mitochondrial volume, respectively.

The dispersion relation of quantum state resulted from the light-mitochondrion coupling system was obtained as follows,

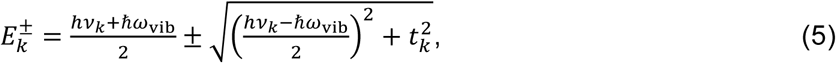

where the plus and minus in “±” indicate the upper and lower branches, respectively; 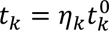, and *η_k_* = Σ*_αγ_**e**_γ_*·***e****_kα_* denotes the direction-matching parameter of light polarization and CH_2_ dipoles. For the energy-active mitochondrion, *η_k_* → 1, and then *t_k_* was estimated to be ∼6.4 kJ mol^-1^ with help of Eq. 4. When *ν_k_* = *ν*_vib_ (i.e., resonance), a superposition state of light and mitochondrion was achieved clearly (Fig. 3e), named a mito-polariton (MP), with the dispersion relation (Eq. 5) reducing to the upper and lower MP levels

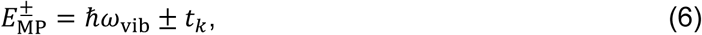

labelled MP+ and MP−, respectively.

#### Correspondence of mitochondrial quantum state with the mitochondrion-related oscillation modes of cells and tissues

To study the relationship of mito-polariton and mitochondrion-related oscillation mode, we compared their properties. Without the coupling of light and mitochondrion, the energy level of mitochondrial lipid CH_2_ vibration excited by an 87-THz photon was at *E*_1_ = *ћω*_vib_ = 34.7 kJ mol^-1^ (Fig. 3e). With the coupling, this excited level was split to two levels: 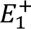 = 28.3 kJ mol^- 1^ for the lower MP−, and 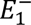 = 41.1 kJ mol^-1^ for the upper MP+, with frequencies of 71 THz and 103 THz, respectively. The level-splitting distance is twice the light-mitochondrion coupling strength *t_k_* (Eq. 6), determined by the structural properties of mitochondria, e.g., the density and order of cristae lipid CH_2_ bonds (Eqs. 4,5). The MP state and level splitting thus will disappear as the mitochondrial structure is destroyed (i.e., *t_k_* → 0), well consistent with our observations that the mitochondrion-related oscillation mode disappears when the HEK-293T cells are ground. Two levels, MP− at 71 THz and MP+ at 103 THz, are expected by the mito-polariton model: MP− can be observed in the spectra due to no vibration modes of biomolecules and water in the 60–80 THz range; whereas MP+ falls in the vibration range (90–110 THz) of peptide amide A and water (Supplementary Fig. 1), thus indistinguishable in our experiments [30,31].

As for the kidney and liver tissues of mice, the lengths of energy-active mitochondria examined through fluorescence microscopy are shown in Fig. 4a (more details in Supplementary Figs. 4,5). Both of them were 2.2 ± 0.4 μm (n = 100 mitochondria), identical to the mitochondrial length of 2.2 ± 0.4 μm in HEK-293T cells. The relation *L*_m_ = 2 × *λ*_m_/2 of standing wave thus exists as well (Fig. 4b upper). Therefore, the MP− level at 71 THz can occur in the mitochondria of kidney and liver tissues like the case of HEK-293T cells.

**Fig. 4.**
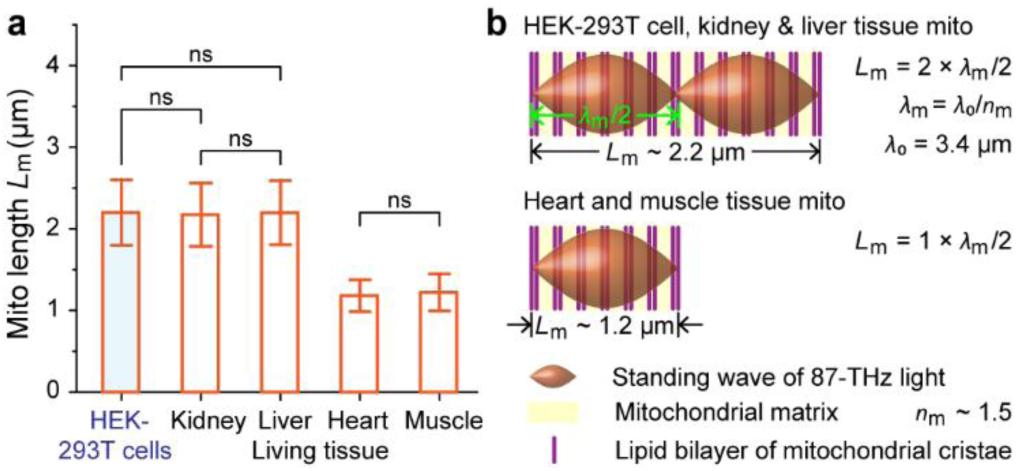
| Relationship of mitochondrial length with the mitochondrial quantum state. **a)** Energy- active mitochondrion lengths of HEK-293T cells and the mouse’s kidney, liver, heart, skeletal muscle tissues. (ns: p > 0.05; t-test, n = 100 mitochondria). **b)** Relationship of mitochondrial length *L*_m_ with the standing wave of 87-THz light in a mitochondrion. The standing-wave condition of light, i.e., *L*_m_ = *m* × *λ*_m_/2, *m* = 1, 2, 3, …, is met and thus the mitochondrion can confine the light, where *λ*_m_ = *λ*_0_/*n*_m_ stands for the light wavelength in mitochondria, *λ*_0_ = 3.4 μm for the vacuum wavelength of 87-THz light and *n*_m_ ∼ 1.5 for the refractive index of mitochondria.

The lengths *L*_m_ of energy-active mitochondria in mouse’s heart and skeletal muscle tissues were both 1.2 ± 0.2 μm (n = 100) (Fig. 4a, more details in Supplementary Figs. 6,7). The wavelength *λ*_m_ of 87-THz light in living mitochondria was ∼2.3 μm, indicating a relation of *L*_m_ = 1 × *λ*_m_/2 (Fig. 4b lower), i.e., the standing-wave condition of *L*_m_ = *m* × *λ*_m_/2 (*m* = 1, 2, 3, …) is still met. The MP− level at 71 THz thus can take place in these cases, corresponding to the mitochondrion-related oscillation modes in the living heart and skeletal muscle tissues we observed.

In all, the coupling of 87-THz light and mitochondrion can cause a quantum superposition state of mito-polariton MP, with a lower level MP− of 71 THz and an upper level MP+ of 103 THz. The MP− corresponds to the mitochondrion-dependent oscillation mode in our experiments; the MP+ falls in the vibration range (90–110 THz) of peptide amide A and water, thus indistinguishable.

### 3. Effect of Mitochondrial Quantum State on the ATP Production of HEK-293T Cells

Mitochondria are primarily responsible for producing ATP in our body. To study the function of mito- polariton, we examined the responses of ATP production in HEK-293T cells to MIR light of 71 THz and 87 THz, respectively (Fig. 5a). MIR lasers of 71.0 THz (4.22 μm in the wavelength) and 87.0 THz (3.45 μm) were employed, with a power density of 10 μW/mm^2^ much lower than the ∼5 mW/mm² applied in optogenetics [41], i.e., a super-weak light.

**Fig. 5.**
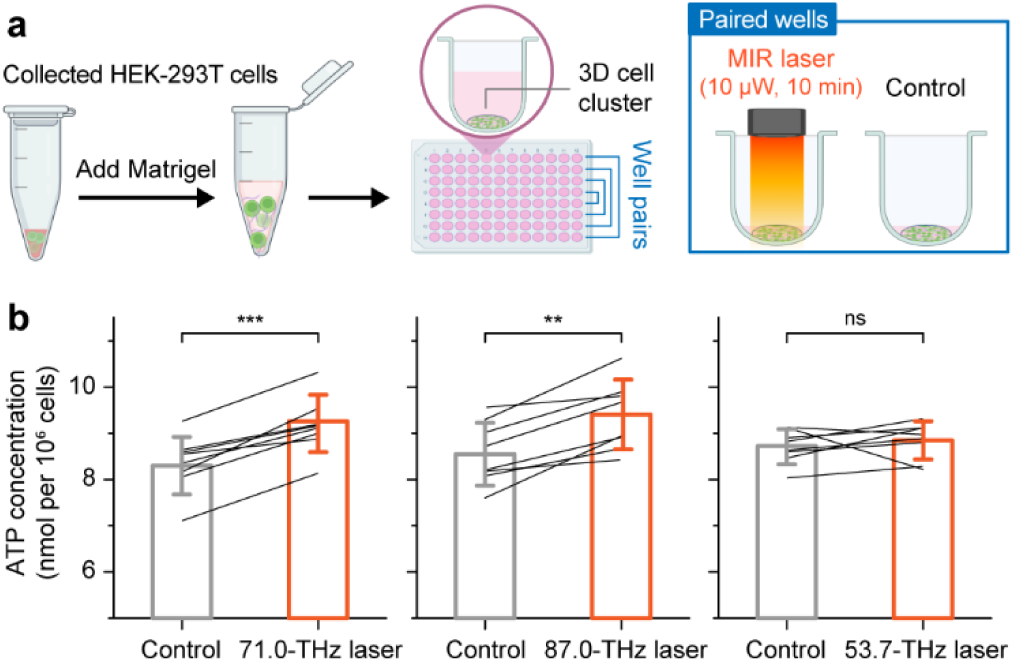
| Modulation of ATP production in living HEK-293T cells by 71.0-THz and 87.0-THz mid- infrared (MIR) light. **a)** Sample preparation and light modulation. An average power density of 10 μW/mm^2^ and an illumination duration of 10 min per sample are applied for the modulation. **b)** ATP production in HEK-293T cells without (gray) and with (orange) the MIR modulation of 71.0 THz (left) and 87.0 THz (middle). Right: A MIR laser of 53.7 THz is further employed as a control. (***: p < 0.001; **: p < 0.01; ns: p > 0.05; t-test, n = 8 samples).

As the results shown in Fig. 5b, compared to control groups (i.e., no MIR modulation), the ATP production under 10-min illumination of the 71.0-THz and 87.0-THz light increased by 10.3% (p < 0.001, n = 8) and 10.1% (p < 0.01, n = 8) with statistical significances, respectively. Both the lasers thus can promote ATP production in the cells. The effect of 71.0-THz light is caused by the resonance with the mitochondrial quantum state of mito-polariton, and that of 87.0-THz light is attributed to the resonant absorption of mitochondrial CH_2_ bonds. A laser of 53.7 THz (5.58 μm in the wavelength) with the identical power density of 10 μW/mm^2^ was further employed as a control. No significant change in ATP production caused by the 10-min illumination of the 53.7-THz light was observed (p > 0.05, n = 8). Together, the mitochondrial quantum state provides an efficient channel to modulate ATP production in living cells.

#### Conclusion

In summary, we demonstrate a quantum state of mitochondria, which can be used to modulate ATP production in living cells. An anomalous 71.0-THz oscillation mode was observed only in living cells and tissues, which is highly determined by mitochondria, and unable to be assigned to any specific molecules. Our experiment data-based analyses and theoretical calculations suggested a quantum superposition state of mitochondrion that is caused by the coupling of light and lipid CH_2_ bonds in functional cristae, and results in the intrinsic CH_2_ vibration mode of 87 THz splitting to the levels of 71 THz and 103 THz. The former corresponds to the anomalous oscillation modes in our experiments; whereas the latter falls in the vibration range (90–110 THz) of peptide amide A and water, thus indistinguishable. Additional experiments revealed that the mitochondrial quantum state can serve as an energy-efficient channel to modulate ATP production. Our findings provide a quantum physics view for understanding living cells. It will be interesting to further explore the physical properties of this unique quantum state, as well as whether the state can act as a channel for energy metabolism, and even information transmission in life.

## Materials and Methods

### MIR spectroscopy of living and dry samples

Fourier Transform Infrared (FTIR) spectroscopy (Thermo Fisher Scientific, Cat# Nicolet 6700) was employed to analyze biological samples under three distinct conditions. For living samples, freshly harvested pellets of HEK-293T cells, mouse tissues, and mitochondria were directly analyzed in the ATR mode (Supplementary Fig. 8). A background of BaF₂ powder suspended in water was used to correct for strong water absorption interference. For thoroughly destroyed dry samples, cell pellets or mouse kidney tissue fragments were completely dehydrated in a 65 °C oven, thoroughly ground with spectroscopic-grade potassium bromide (KBr) at a 1:200 ratio, and pressed into pellets for the transmission mode under N₂ purge (Supplementary Fig. 9). For all the measurements, instrument-specific background spectra were acquired prior to sample analysis. Seeing more in Supplementary Information.

### 3D confocal imaging and structural analysis of MMP-higher mitochondria

Living HEK-293T cells and fresh 40-μm tissue sections of mouse were stained in 35-mm glass-bottom dishes. For immediate imaging, samples were incubated with the mitochondrial membrane potential probe JC-1 (Thermo Fisher Scientific, Cat# T3168, 2.5 μg/mL) and the nuclear dye Hoechst 33342 (Beyotime, Cat#C1027, 1×) in medium at 37 °C under 5% CO₂ for 30 minutes (cells) or 2 to 4 hours (tissues).

3D fluorescence imaging was performed using an inverted confocal microscope (Leica DMi8) equipped with a 63× oil-immersion objective. Z-stacks were acquired with step sizes of 0.15 μm (cells) or 0.25 μm (tissues). For analysis, 3D reconstructions were generated and background-corrected using LAS X software, then segmented into 5-μm-thick slices along the y-axis. Fluorescence intensity and morphological parameters of higher-MMP mitochondria were quantified using ImageJ.

### MIR light source, cell illumination and ATP assay

Tunable quantum cascade lasers (HUBNER Photonics, Cobolt Odin™ Series, 3450 nm; MapleFire Solution, 4220nm and 5580 nm) served as the MIR sources, with the output coupled into a hollow-core MIR fiber (Guiding Photonics, HF500MW -SMA-Gn). The output power at the fiber tip was 10 µW.

To match the ∼1 mm laser spot size, a 3D culture system was established (Supplementary Fig. 10). HEK-293T cells were resuspended and mixed with an equal volume of pre-melted Matrigel (Corning, Cat #356237) on ice. A 3 µL aliquot of the mixture (∼60,000 cells) was dispensed into the center of wells in a U-bottom 96-well plate and polymerized at 37°C for 30 minutes to form cell clusters approximately 1 mm in diameter. Each well was then supplemented with 150 µL of culture medium and incubated at 37°C under 5% CO₂ for 6–12 hours prior to experimentation.

For illumination, optical fibers delivered light into a portable incubator maintained at 37°C and 5% CO₂ (Supplementary Fig. 11). For each experiment, two symmetrically positioned wells within the same column were selected. After removing the old medium and adding only 5 µL of fresh medium, one well received MIR illumination of 10 min, with its paired well as an untreated control.

Following illumination, cells were lysed with ice-cold ATP lysis buffer from a luciferase-based ATP assay kit (Beyotime, Cat#S0027). The lysates were centrifuged, and the supernatants were transferred to a white 96-well plate. ATP levels were quantified using the ATP assay working solution and a microplate reader. The data were normalized to cell number and compared between groups using a paired Student’s t-test (n = 8). Seeing more in Supplementary Information.

## Acknowledgments.

This work was supported by the National Natural Science Foundation of China (T2394532, T2241002) and the National Key R&D Program of China (2021YFA1200404).

## Author Contributions

B.S. and Z.G. conceived the research; B.S., Z.G and Y.Y. designed the experiments, Y.Y. performed the experiments; B.S. designed and performed the quantum mechanics calculations. All authors analyzed the results. B.S., Z.G and Y.Y. wrote the manuscript.

---

**Supplementary Fig. 1.**
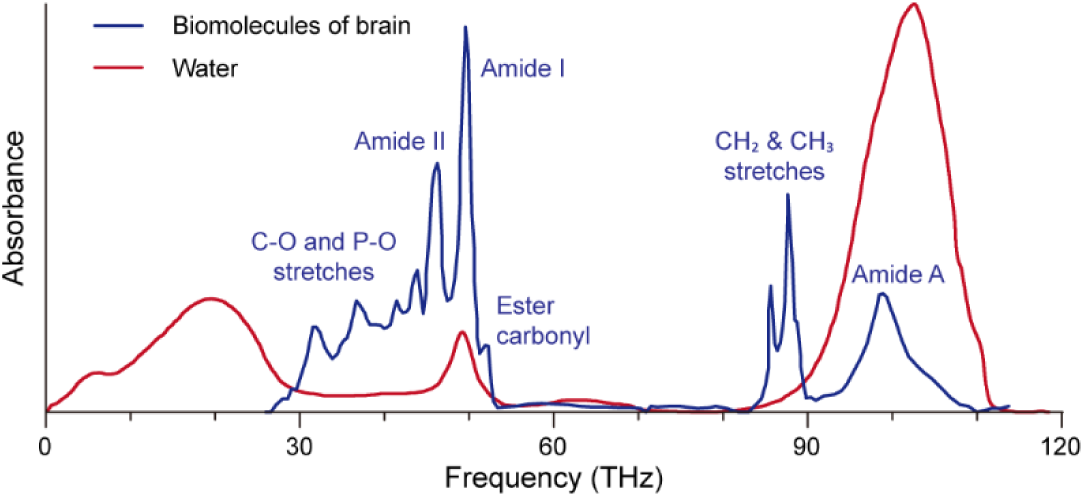
| Infrared absorption spectra of biomolecules (blue) and water (red). Water is usually considered as a necessary part of biological systems. Reproduced with permissions from Ref. 30 Copyright 2011 Elsevier and from Ref. 31 Copyright 2012 Elsevier, respectively.

**Supplementary Fig. 2.**
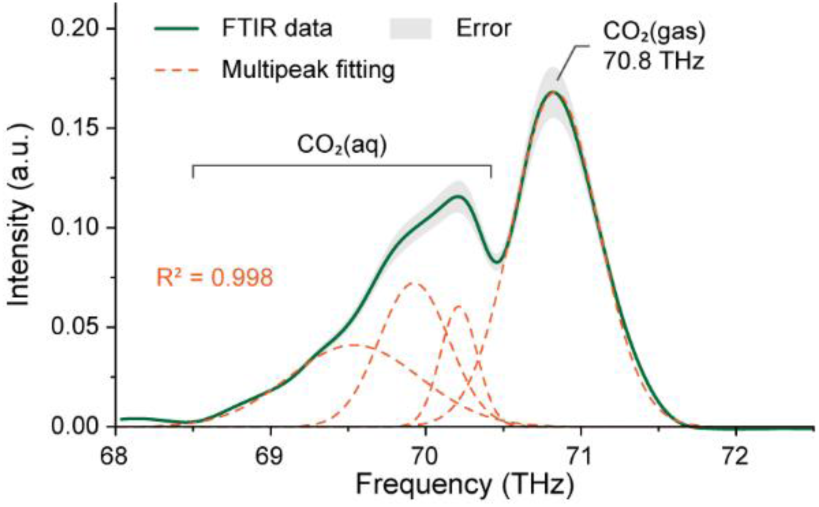
| CO_2_ vibration modes and their multipeak analysis. The measurement is conducted by FTIR-ATR. The multipeak analysis with the determination coefficient R^2^ > 0.995 shows that a peak is clearly located at 70.8 THz, which can be assigned to CO_2_(gas) vibration, while there are at least three peaks in the range of 68.5–70.4 THz, which can be assigned to CO_2_(aq) vibration.

**Supplementary Fig. 3.**
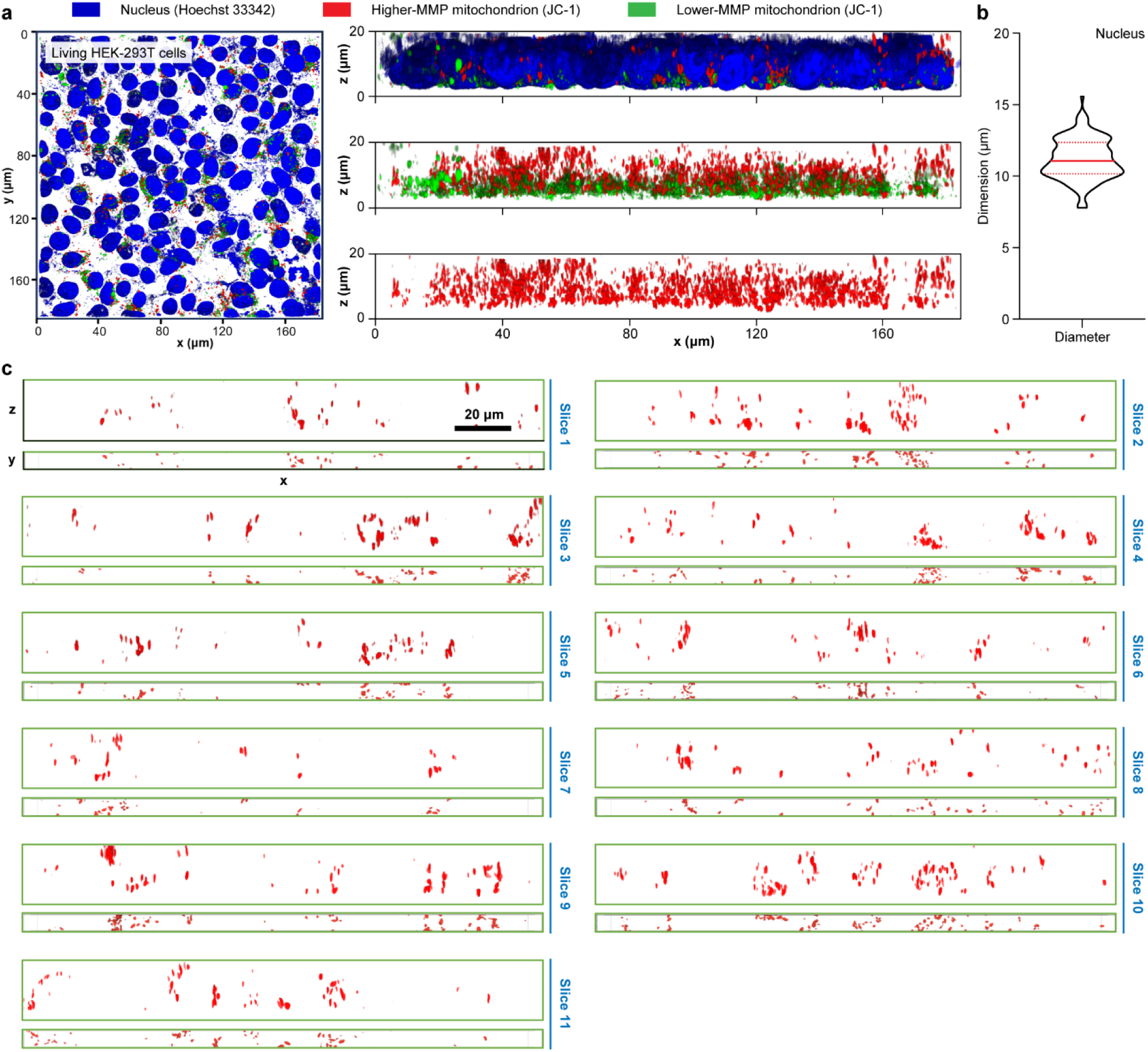
| Confocal fluorescence microscopy imaging analyses of the nuclei and mitochondria in living HEK-293T cells. **a)** Fluorescence microscopy image. The blue indicates the nucleus (Hoechst 33342 stained); the red and green mean the mitochondria with higher and lower MMP (JC-1 stained), respectively. Left: x-y view. Right: x-z view. **b)** Nuclear diameter. The solid and dashed red lines denote the average and standard deviation, respectively. **c)** All 5-μm x-z plane slices in the x-z and x-y views for the statistics of higher-MMP mitochondrial dimensions.

**Supplementary Fig. 4.**
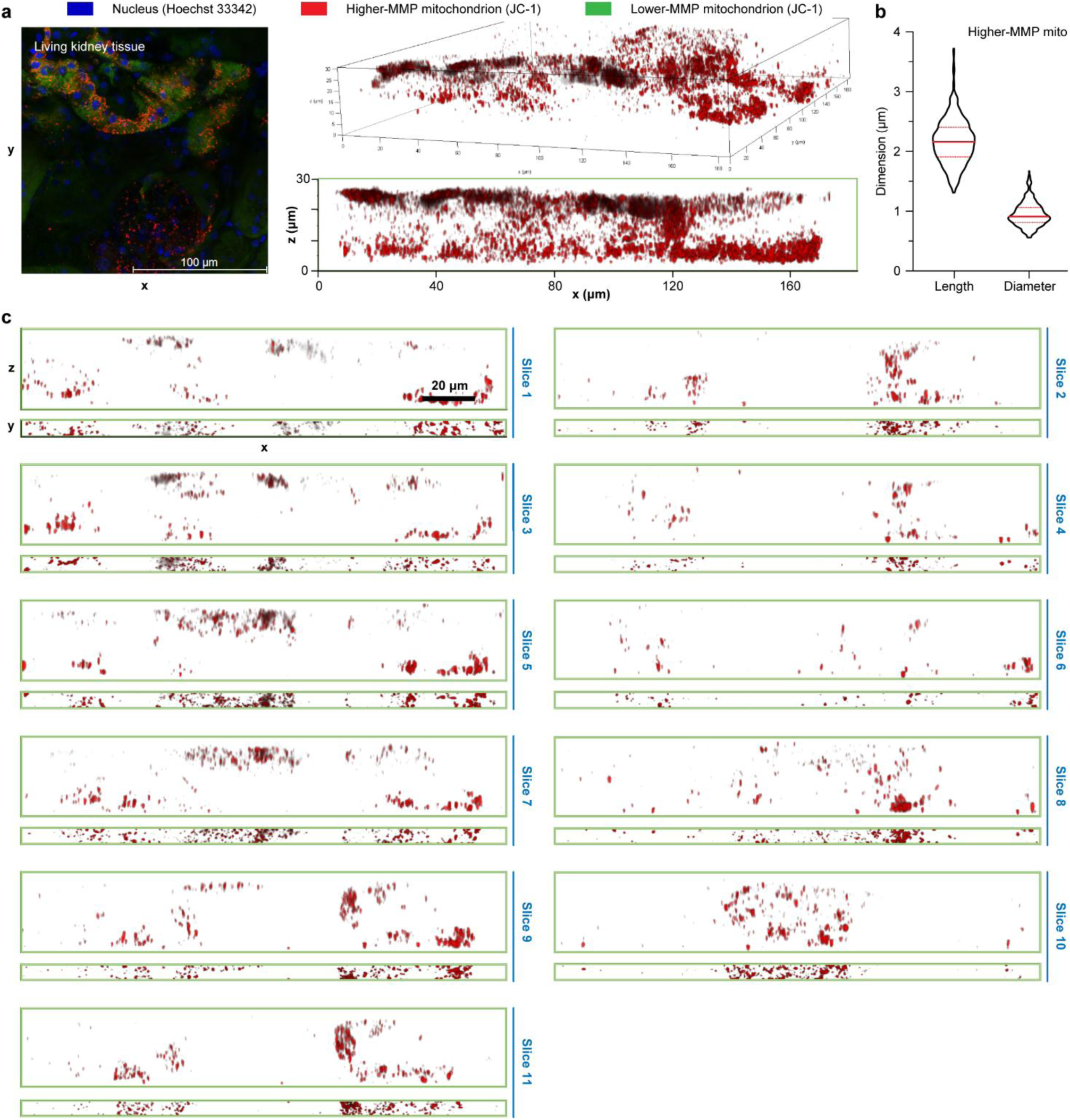
| Confocal fluorescence microscopy imaging analyses on the length and width of higher-MMP mitochondria in the living kidney tissue of mice. **a)** Fluorescence microscopy image. The blue indicates the nucleus (Hoechst 33342 stained); the red and green represent the mitochondria with and without higher MMP (JC-1 stained), respectively. **b)** Length and maximum-diameter of the complete mitochondria with higher MMP. **c)** All 5-μm x-z plane slices in the x-z and x-y views for the statistics of higher-MMP mitochondrial dimensions.

**Supplementary Fig. 5.**
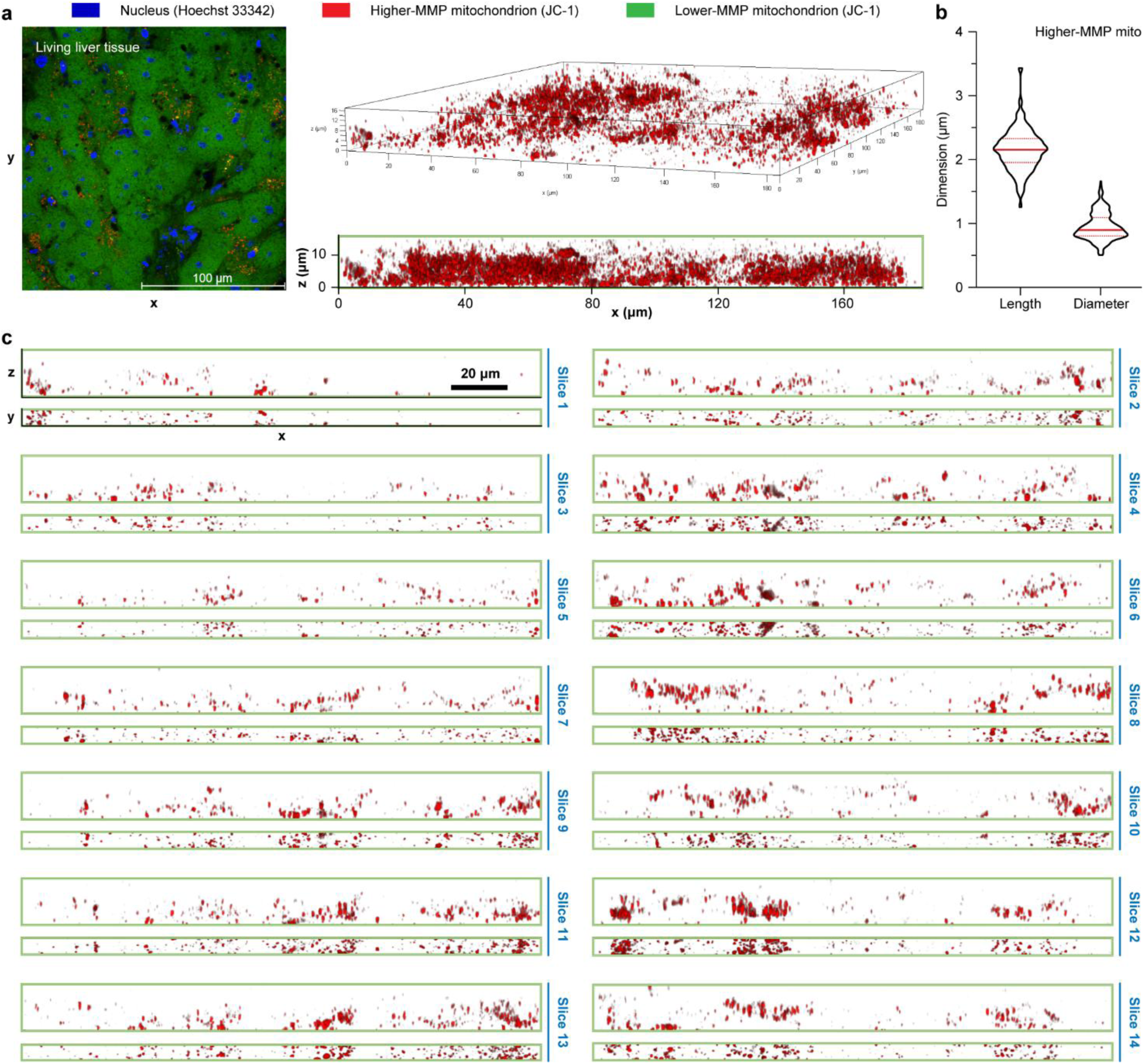
| Confocal fluorescence microscopy imaging analyses on the length and width of higher-MMP mitochondria in the living liver tissue of mice. **a)** Fluorescence microscopy image. The blue indicates the nucleus (Hoechst 33342 stained); the red and green mean the mitochondria with and without higher MMP (JC-1 stained), respectively. **b)** Length and maximum- diameter of the complete mitochondria with higher MMP. The solid and dashed red lines denote the average and standard deviation, respectively. **c)** All 5-μm x-z plane slices in the x-z and x-y views for statistics of higher-MMP mitochondrial dimensions.

**Supplementary Fig. 6.**
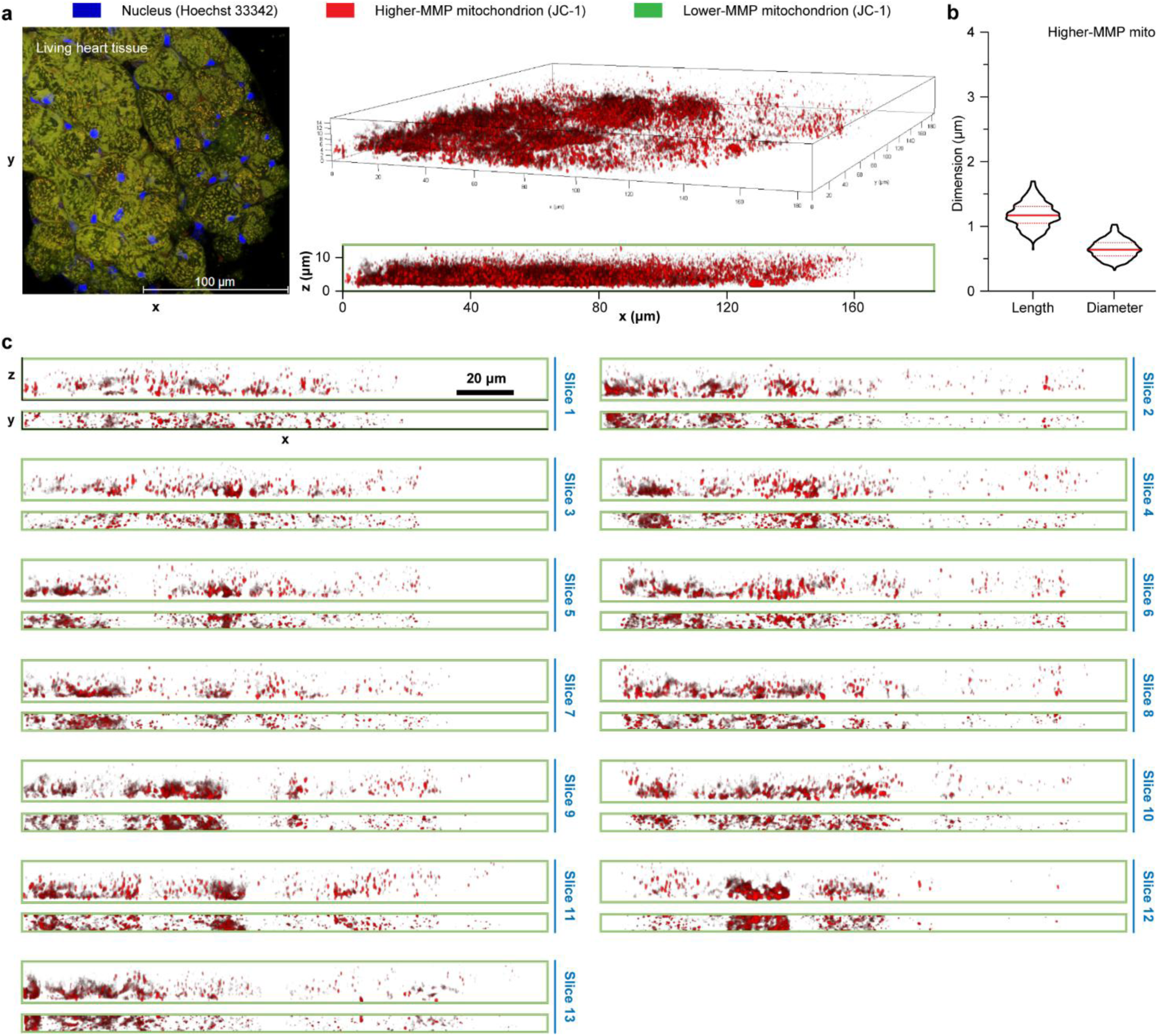
| Confocal fluorescence microscopy imaging analyses on the length and width of higher-MMP mitochondria in the living heart tissue of mice. **a)** Fluorescence microscopy image. The blue indicates the nucleus (Hoechst 33342 stained); the red and green mean the mitochondria with and without higher MMP (JC-1 stained), respectively. **b)** Length and maximum- diameter of the complete mitochondria with higher MMP. The solid and dashed red lines denote the average and standard deviation, respectively. **c)** All 5-μm x-z plane slices in the x-z and x-y views for the statistics of higher-MMP mitochondrial dimensions.

**Supplementary Fig. 7.**
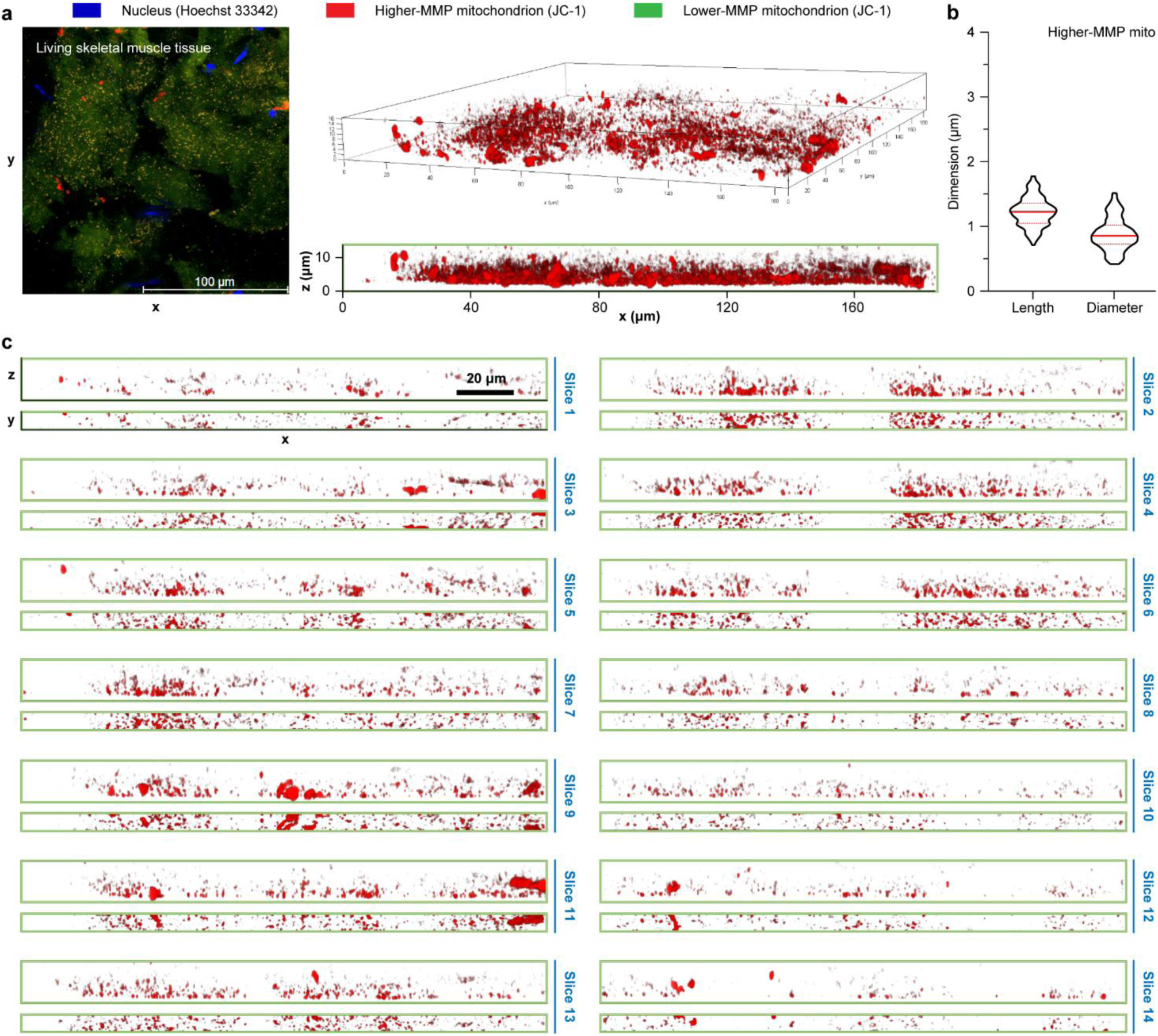
| Confocal fluorescence microscopy imaging analyses on the length and width of higher-MMP mitochondria in the living skeletal muscle tissue of mice. **a)** Fluorescence microscopy image. The blue indicates the nucleus (Hoechst 33342 stained); the red and green represent the mitochondria with and without higher MMP (JC-1 stained), respectively. **b)** Length and maximum-diameter of the complete mitochondria with higher MMP. The solid and dashed red lines mean the average and standard deviation, respectively. **c)** All 5-μm x-z plane slices in the x-z and x-y views for the statistics of higher-MMP mitochondrial dimensions.

**Supplementary Fig. 8.**
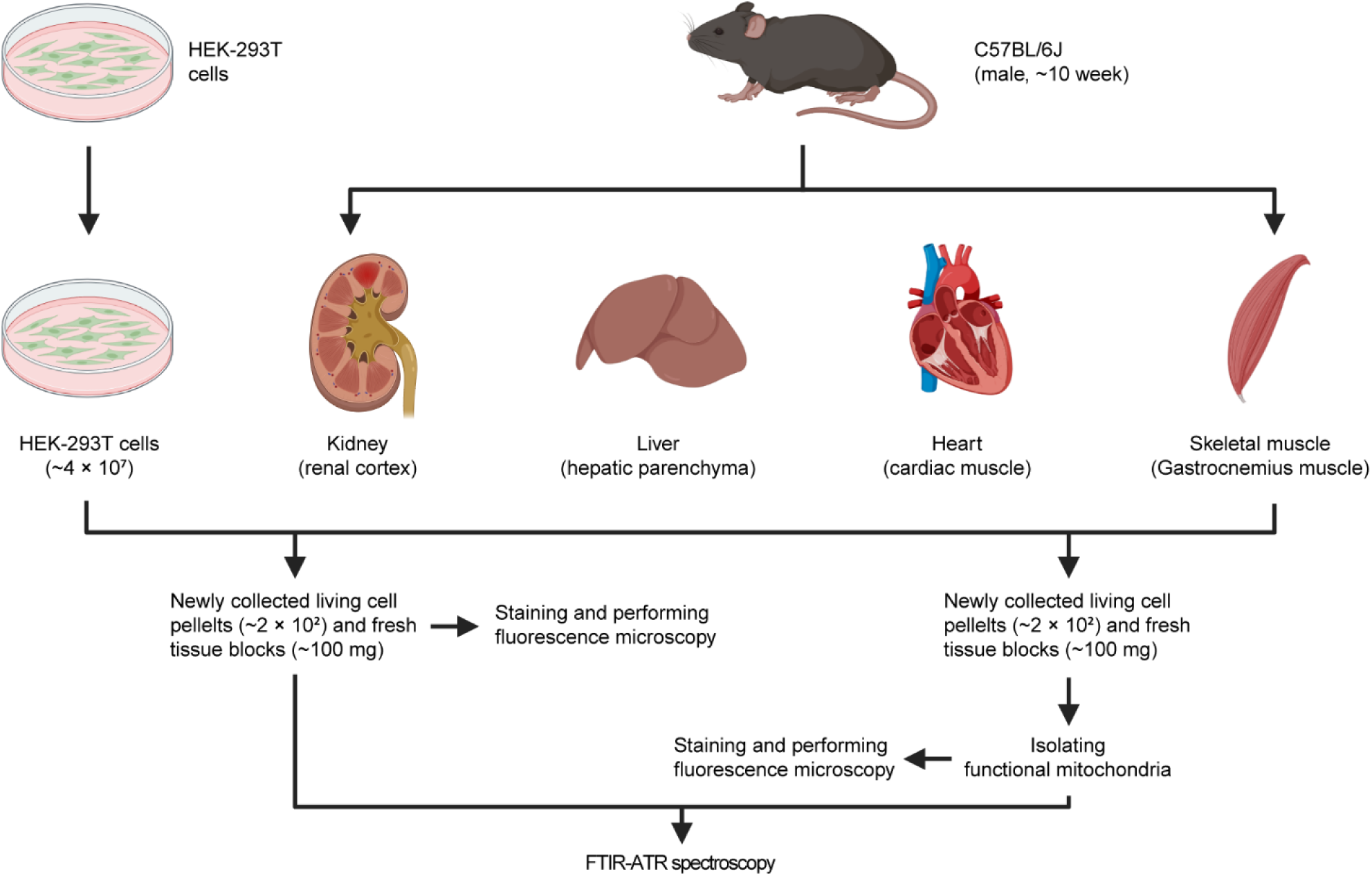
| Sample preparation and experiment details for the FTIR spectroscopy of living HEK-293T cells, mouse tissues and mitochondria.

**Supplementary Fig. 9.**
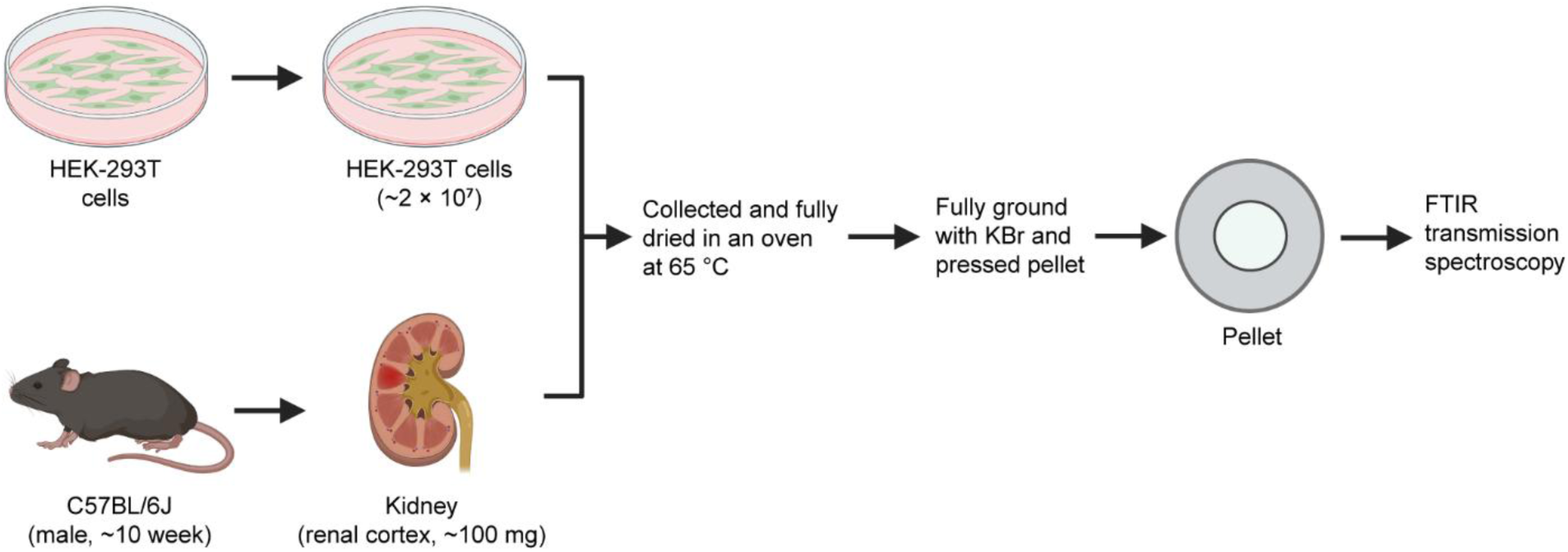
| Sample preparation and experiment details for the FTIR spectroscopy of thoroughly ground dry HEK-293T cells and mouse kidney tissue.

**Supplementary Fig. 10.**
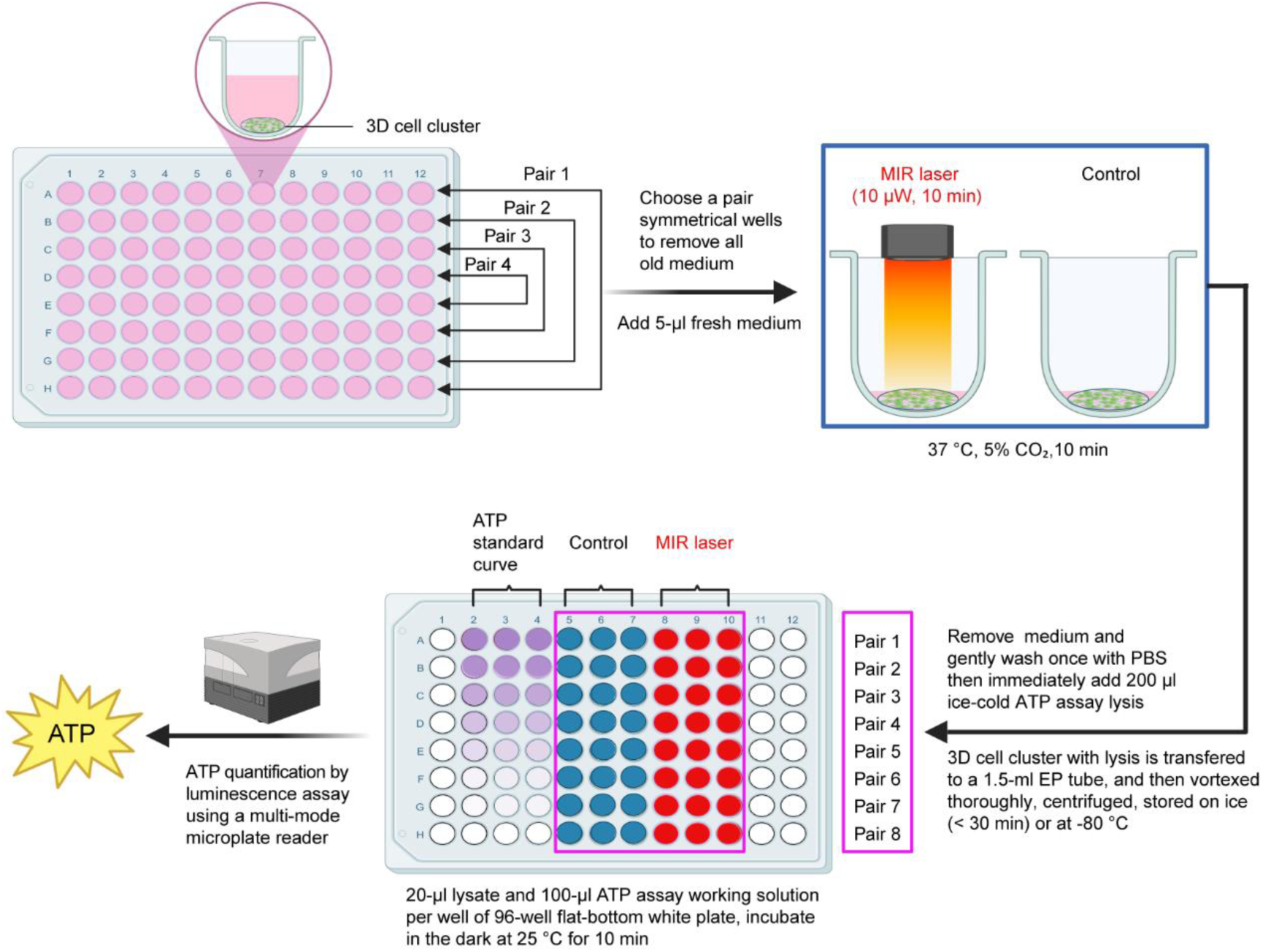
| Experimental details of MIR modulation and ATP detection on HEK- 293T cells. Two symmetrical wells were selected as a pair each time (to ensure consistency in growth conditions between pairs as much as possible). After removing the old culture medium, 5 μl of fresh culture medium was added. The MIR treatment was then immediately performed according to the protocol shown in the figure. Each treatment was replicated across 8 experimental groups. Each sample was treated with 200 μl of ATP assay lysis and thoroughly mixed. Subsequently, the ATP detection protocol was followed: three replicate wells were set up for each sample, with 20 μl of lysate added to each well. In the ATP standard curve wells, 20 μl of ATP standard dilutions were added. After that, 100 μl of ATP assay working solution was rapidly added to all wells, followed by incubation at room temperature in the dark for 10 minutes. Fluorescence luminescence signals were quantified using a multifunctional microplate reader. Finally, the ATP concentrations for each well were calculated based on the standard curve, and the average ATP production per sample was determined by normalizing to the original cell quantity.

**Supplementary Fig. 11.**
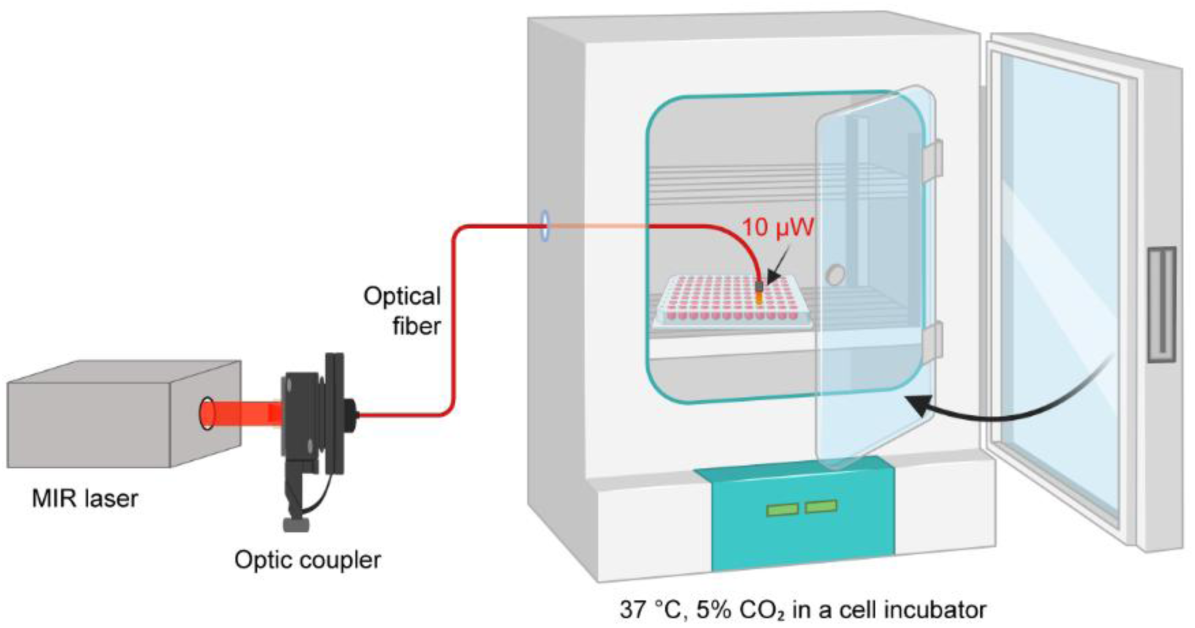
| Optical path diagram of MIR modulation.

## References

1. M. E. Raichle, D. A. Gusnard. Appraising the brain’s energy budget. Proc. Natl. Acad. Sci. U.S.A. 99, 10237–10239 (2002).

2. M.-D. Cernescu, M. Butnariu. The body’s energy requirement; general considerations. J. N. Food Sci. Tech. 5, 1–3 (2024).

3. S. Hameroff, R. Penrose. Orchestrated reduction of quantum coherence in brain microtubules: a model for consciousness. Math. Comput. Simulat. 40, 453–480 (1996).

4. Y. Wang et al. A physical derivation of high-flux ion transport in biological channel via quantum ion coherence. Nat. Commun. 15, 7189 (2024).

5. G. S. Engel et al. Evidence for wavelike energy transfer through quantum coherence in photosynthetic systems. Nature 446, 782–786 (2007).

6. H. Lee et al. Coherence dynamics in photosynthesis: protein protection of excitonic coherence. Science 316, 1462–1465 (2007).

7. Y.-C. Chen et al. Resonant confinement of an excitonic polariton and ultraefficient light harvest in artificial photosynthesis. Phys. Rev. Lett. 122, 257402 (2019).

8. S. Pedalino, B. E. Ramírez-Galindo, R. Ferstl, K. Hornberger, M. Arndt, S. Gerlich. Probing quantum mechanics with nanoparticle matter-wave interferometry. Nature 649, 866–870 (2026).

9. X. Liu et al. Nonthermal and reversible control of neuronal signaling and behavior by midinfrared stimulation. Proc. Natl. Acad. Sci. U.S.A. 118, e2015685118 (2021).

10. J. Zhang et al. Non-invasive, opsin-free mid-infrared modulation activates cortical neurons and accelerates associative learning. Nat. Commun. 12, 2730 (2021).

11. Y. Yang et al. AuNP-modulated qPCR: an optimized system for detecting MIR biophotons released in DNA replication. Chem. Eur. J. 29 e202203513 (2023).

12. N. Li et al. Demonstration of biophoton-driven DNA replication via gold nanoparticle-distance modulated yield oscillation. Nano Res. 14, 40–45 (2021).

13. C. Zhang et al. Driving DNA origami assembly with a terahertz wave. Nano Lett. 22, 468–475 (2022).

14. B. Song, Y. Shu. Cell vibron polariton resonantly self-confined in the myelin sheath of nerve. Nano Res. 13, 38– 44 (2020).

15. D. Peng et al. Mid-infrared photons released by NAD^+^ reduction in the tricarboxylic acid cycle of myelinated neuron. Neurosci. Bull. 39, 1146–1150 (2023).

16. Z. Liu et al. Entangled biphoton generation in the myelin sheath. *Phys*. Rev. E 110, 024402 (2024).

17. R. M. Schwartz, M. O. Dayhoff. Origins of prokaryotes, eukaryotes, mitochondria, and chloroplasts. Science 199, 395–403 (1978).

18. J. Vosseberg et al. The emerging view on the origin and early evolution of eukaryotic cells. Nature 633, 295– 305 (2024).

19. C. Lopez-Otin et al. Hallmarks of aging: an expanding universe. Cell 186, 243–278 (2023).

20. D. C. Wallace. Mitochondria and cancer. Nat. Rev. Cancer 12, 685–698 (2012).

21. M. P. Murphy, L. A. J. O’Neill. A break in mitochondrial endosymbiosis as a basis for inflammatory diseases. Nature 626, 271–279 (2024).

22. P. Gonzalez-Rodriguez et al. Disruption of mitochondrial complex I induces progressive parkinsonism. Nature 599, 650–656 (2021).

23. C. F. Matta, L. Massa. Energy equivalence of information in the mitochondrion and the thermodynamic efficiency of ATP synthase. Biochem. 54, 5376–5378 (2015).

24. M. Wikström, R. Springett. Thermodynamic efficiency, reversibility, and degree of coupling in energy conservation by the mitochondrial respiratory chain. *Commun*. Biol. 3, 451 (2020).

25. C. C. Moser et al. Electron tunneling chains of mitochondria. Biochem. Biophys. Acta 1757, 1096–1109 (2006).

26. H. Xin et al. Quantum biological tunnel junction for electron transfer imaging in live cells. Nat. Commun. 10, 3245 (2019).

27. S. de Vries et al. Electron tunneling rates in respiratory complex I are tuned for efficient energy conversion. Angew. Chem. Int. Ed. Engl. 54, 2844–2848 (2015).

28. P. A. M. Dirac, The Principles of Quantum Mechanics (Oxford: At the Clarendon Press, 1930).

29. J. Fahrenfort. Attenuated total reflection: A new principle for the production of useful infra-red reflection spectra of organic compounds. Spectrochim. Acta 17, 698–709 (1961).

30. Y. Maréchal. The molecular structure of liquid water delivered by absorption spectroscopy in the whole IR region completed with thermodynamics data. J. Mol. Struct. 1004, 146–155 (2011).

31. S. Caine et al. The application of Fourier transform infrared microspectroscopy for the study of diseased central nervous system tissue. Neuroimage 59, 3624–3640 (2012).

32. T. Schädle, B. Pejcic, B. Mizaikoff. Monitoring dissolved carbon dioxide and methane in brine environments at high pressure using IR-ATR spectroscopy. Anal. Methods 8, 756–762 (2016).

33. J. W. Ellwart, P. Dormer. Vitality measurement using spectrum shift in Hoechst-33342 stained cells. Cytometry 11, 239–243 (1990).

34. S. T. Smiley et al. Intracellular heterogeneity in mitochondrial membrane potentials revealed by a J-aggregate-forming lipophilic cation JC-1. Proc. Natl. Acad. Sci. U.S.A. 88, 3671–3675 (1991).

35. M. Lherbette et al. Atomic force microscopy micro-rheology reveals large structural inhomogeneities in single cell-nuclei. Sci. Rep. 7, 8116 (2017).

36. R. Flindt. Amazing Numbers in Biology (Springer-Verlag Berlin Heidelberg, 2006), pp. 254.

37. T. Liu et al. Multi-color live-cell STED nanoscopy of mitochondria with a gentle inner membrane stain. Proc. Natl. Acad. Sci. U.S.A. 119, e2215799119 (2022).

38. N. Kucerka et al. Structure of fully hydrated fluid phase lipid bilayers with monounsaturated chains. J. Membr. Biol. 208, 193–202 (2005).

39. S. Cogliati, J. A. Enriquez, L. Scorrano. Mitochondrial cristae: where beauty meets functionality. Trends Biochem. Sci. 41, 261–273 (2016).

40. J. Kwon et al. Label-free nanoscale optical metrology on myelinated axons in vivo. Nat. Commun. 8, 1837 (2017).

41. E. S. Boyden et al. Millisecond-timescale, genetically targeted optical control of neural activity. Nat. Neurosci. 8, 1263–1268 (2005).

